# Taxonomic, temporal, and spatial variation in the dynamics of High-Arctic arthropod populations

**DOI:** 10.1101/2020.04.27.052795

**Authors:** Toke T. Høye, Sarah Loboda, Amanda M. Koltz, Mark A. K. Gillespie, Joseph J. Bowden, Niels M. Schmidt

## Abstract

Time-series data on arthropod populations are critical for understanding the magnitude, direction, and drivers of abundance changes. However, most arthropod monitoring programs are short-lived and limited in taxonomic resolution and spatial extent. Consequently, variation in population dynamics among taxa and habitats remains poorly understood. Monitoring data from the Arctic are particularly underrepresented, yet important to assessments of species abundance changes because many anthropogenic drivers of change that are present in other regions are absent in polar regions. Here, we utilise 24 years of abundance data from Zackenberg in High-Arctic Greenland, which is the longest running Arctic arthropod monitoring program, to study temporal trends in abundance. Despite a strong warming signal in air temperature, we only find evidence of weak temporal trends in arthropod abundances across most taxa. These trends are more pronounced in the most recent decade, with change point analyses suggesting distinct non-linear dynamics within some functional groups such as predators and detritivores. Although the abundances of many taxa were correlated, we detected both positive and negative correlations, suggesting that multiple processes are affecting arthropod populations even in this relatively simple Arctic food web. Finally, we found clear differences among species within single families of arthropods, indicating that an apparent lack of change in abundance at broader taxonomic or functional levels could mask substantial species-specific trends. Our results reiterate the need for more basic research into the life-history, ecology, and adaptation of arthropod species to better understand their sensitivity to global changes.

**Significance statement:** Terrestrial arthropods, including insects and spiders, serve critical ecosystem functions and are excellent indicators of environmental change due to their physiology, short generation time, and abundance. The Arctic, with its rapid climate change and limited direct anthropogenic impact, is ideal for examining arthropod population dynamics. We use the most comprehensive, standardized dataset available on Arctic arthropods to evaluate the variability in population dynamics for the most common arthropod groups at various taxonomic levels across 24 years. Our results highlight that temporal trends of arthropod populations seem less directional in the Arctic than in temperate regions. Although abundances of some arthropod taxa are declining, particularly in recent decades, population trends still display high variation among time periods, taxa, and habitats.

## INTRODUCTION

The abundance and diversity of terrestrial arthropods are under threat from several anthropogenic pressures (e.g. 1, 2-8). Long-term monitoring data indicate that abundances of many arthropod species are declining, as shown in a meta-analysis study that documented declines in abundances of >450 invertebrate species globally (9). However, each assessment of a decline has its various strengths and weaknesses. Some are based on biological records requiring corrections for sampling effort (1, 4, 6, 7), while others only provide a comparison of discrete points in time (4, 10). These, and other shortfalls have strengthened calls for more standardized, long-term biological monitoring (11-14), including in areas less dominated by humans where drivers other than agricultural intensity can be investigated (15-17).

Previous work on species declines has suggested that the primary pressures on arthropod populations and communities in recent decades include habitat fragmentation, habitat loss, and land-use intensification (2, 18), though climate change will likely increase in importance as warming continues, even at low latitudes (19). Teasing apart the effects of land-use change and climate on arthropods, however, remains a major challenge (17). For example, while Hallmann, *et al.* (3) found a mid-summer decline in flying insect biomass of 82% over 27 years in protected areas in Germany, this pattern could not be attributed to landscape or climatic factors. Similarly, the most apparent and recent evidence of declines in arthropod biomass, abundance, and richness from standardized sampling could not confirm hypotheses about the impact of local land-use intensity (2). The impacts of individual drivers associated with land-use intensity, such as pesticide use, are particularly difficult to separate without a careful experimental design incorporated into monitoring systems (17, 20), and particularly so over the long-term.

Arctic regions are useful to study the impacts of climate change because many direct anthropogenic disturbances are absent (21), and because these regions are warming rapidly (22). While long term data are generally scarce in the Arctic (15), the arthropod monitoring program at Zackenberg, North-East Greenland, has been collecting standardised data since 1996, representing the longest running program in the Arctic (23). These data offer a rare opportunity to detect empirical change in a broad array of taxa, not least because of the availability of samples across taxonomic groups in different habitats. Many long-term monitoring programs that include arthropods often only report trends at coarse taxonomic resolution or focus on single habitat types (3, 8, 24, 25). This can mask important variation, which can be teased apart only with analyses of long-term population dynamics of arthropods from different habitats and taxa (e.g. 26, 27, 28). Our previous work on spiders (Araneae) and a single family of flies (Diptera) has shown that habitat type can play an important role in the strength of species abundance trends (16, 28, 29). Previously, we also found evidence of differential long-term changes across invertebrate orders, altering the entire community composition (30). Therefore, there is an urgent need for an improved understanding of the spatial and taxonomic variation in population dynamics of arthropods. The Zackenberg dataset is unique in addressing these issues by including sampling across taxonomic groups, in different habitat types, and across multiple decades.

In this study, we use the Zackenberg dataset consisting of > 1 million individuals collected and counted over the last 24 years to address some of the above issues with long-term data reporting, while also improving our understanding of long-term change in a community of terrestrial and semi-aquatic arthropods under rapid climate change. Our specific objectives are 1) to assess and compare the temporal dynamics in climate and arthropod abundance across taxa, habitats, and time periods by estimating linear trends as well as non-linear dynamics, 2) to quantify shared and opposing population dynamics among arthropod taxa using cross-correlations, and finally, 3) to assess the credibility of using trends in broader taxonomic groups as indicators of single species by comparing species-level and family-level trends in abundance using available data from a subset of 18 years. Since no direct anthropogenic pressures occur at this site, our baseline assumption is that significant changes in arthropod abundances are a result of direct (e.g., of temperature on physiology) and indirect (e.g., via species interactions) effects of climate change.

## RESULTS

We found only limited evidence for long-term abundance trends in insect, spider, and micro-arthropod populations at Zackenberg from a total of 1,006,848 individuals collected across the entire 24-year time period (Figure 1a). An explanation of the taxonomic affiliation and broad functional group of the taxa included is presented in Table S1. When the study period was broken up into shorter decadal time-windows, we still found few significant trends (n=3) in the first decade (Figure 1b), but more significant trends in the central (n=9; Figure 1c), and last (n=13) decades (Figure 1d). Trends in abundance were generally a mix of positive and negative, although in the last decade more than two thirds of the significant trends were negative (Figure 1).

**Figure 1.**
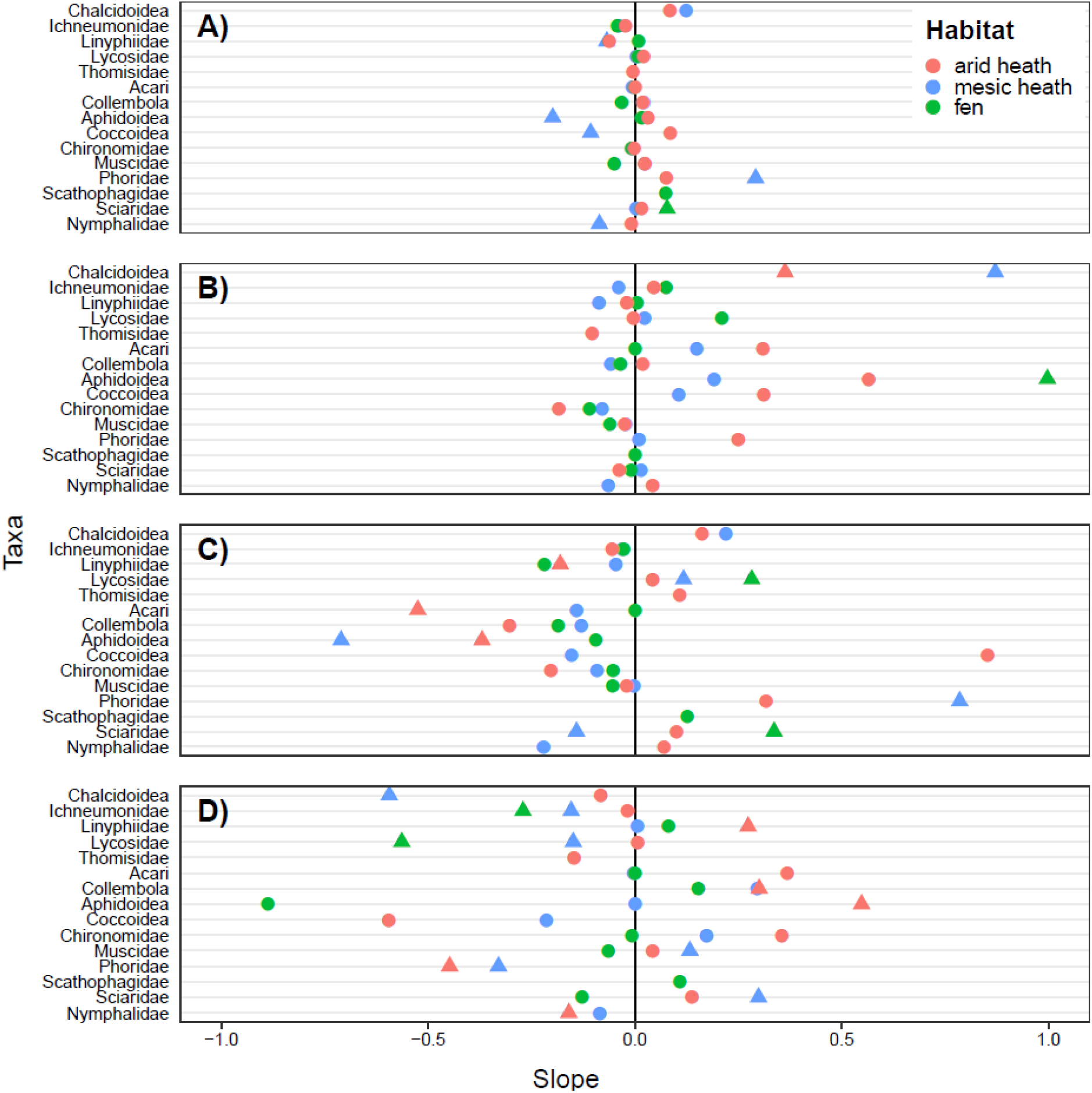
Trend analysis at family or higher taxonomic level grouped according to broad functional groups as outlined in Table S1 a) across the entire study period, b) during the first decade (1996–2005), c) during the middle decade (2002–2011), and d) during the last decade (2009–2018) for each habitat. Temporal trends are estimated with the modified Mann-Kendall non-parametric trend analysis and significant trends at p < 0.05 are indicated by triangles.

We used principal component analysis (PCA) to summarize several climate variables from Zackenberg of relevance to the terrestrial arthropod community. The PCA with varimax rotation revealed that 69% of the variation in our climatic variables was explained by the first three axes of the PCA (see Table S2 for relative contributions of each of the climatic variables to the derived PCs). Higher values of PC1 indicate warmer previous fall temperatures and higher summer precipitation, of PC2 indicate warmer previous summer and shorter winter duration, and of PC3 warmer winter and spring temperatures (Figure 2). We then applied change point analysis to these PCs to detect whether there was evidence of abrupt and linear climatic change over the study period, and to visually assess whether these changes corresponded to changes in arthropod abundances. We found significant change points in all three PCs (Figure 2, Table S3) as well as in several individual climate variables (Figure S1). The change point analysis of arthropod abundance also revealed distinct commonalities in the dynamics of functionally related taxa. Specifically, the highest abundances of predators and parasitoids were detected during the central part of the study period. Conversely, decomposers experienced reduced abundances in the central part of the study period and relatively high abundances in the earlier and more recent years. Within the Diptera, which are diverse in their ecosystem function, we found contrasting and complex dynamics between groups. Herbivores and pollinators were stable throughout the study period in arid heath and generally rare in the fen. In mesic heath, they were generally declining with a distinct drop halfway through the study period. Detailed results from the change point analyses of individual taxa from each habitat are presented in Table S4 and analysis of linear trends for those cases where no break points were identified are presented in Table S5.

**Figure 2.**
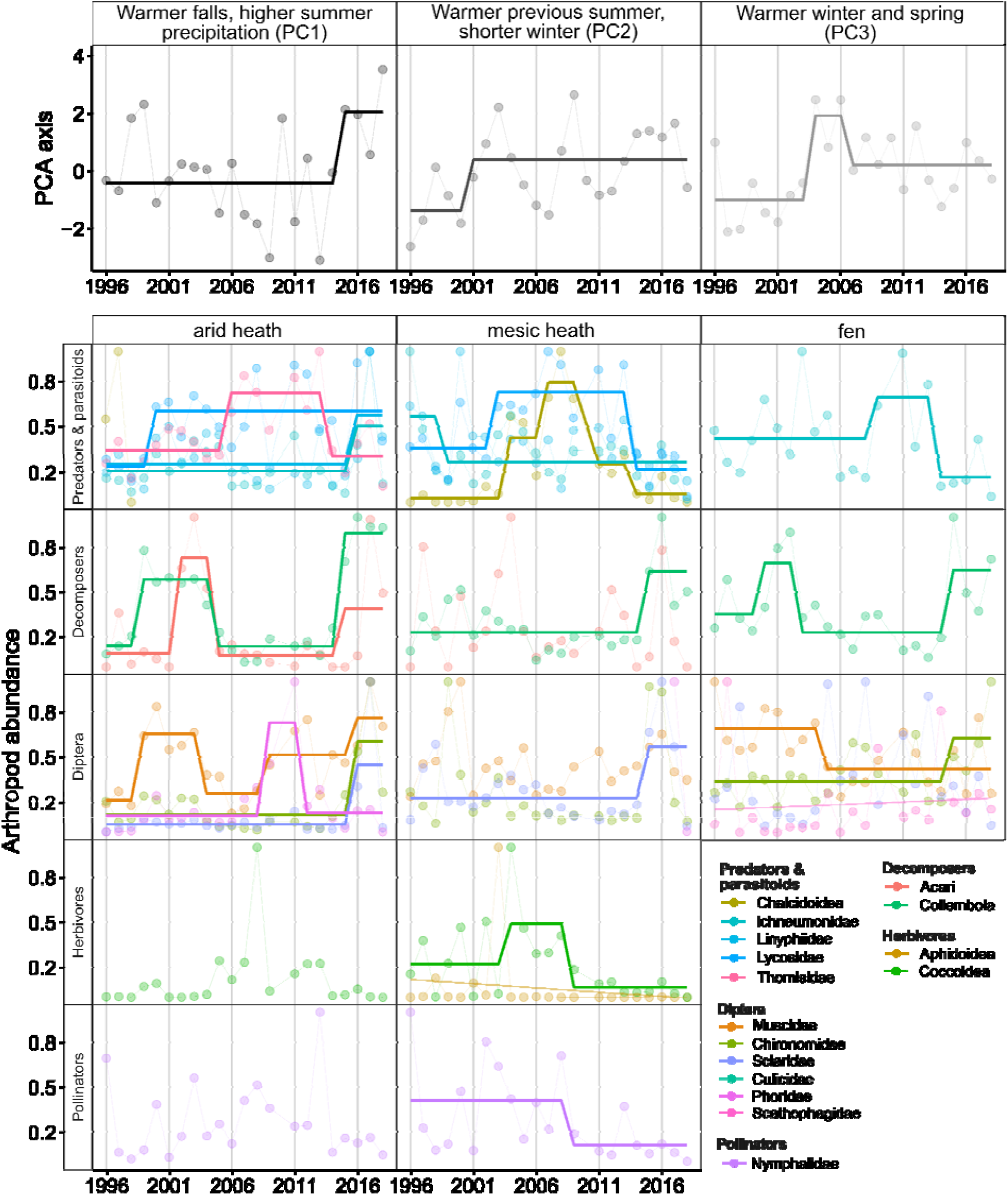
Inter-annual variation and change point analysis of key climate variables of relevance to arthropods as summarized in the three PCA axes and arthropod abundance at the family and higher taxonomic levels. Arthropod taxa are grouped into broad functional groups as outlined in Table S1. Solid lines are drawn where the change point analysis was significant (see detailed results in Tables S3, S4, and S5). Arthropod data are not available from 2010.

Shared population dynamics among arthropod taxa would indicate that they are driven by similar environmental factors or biotic feedbacks. Without knowing the exact regulating factors for the species belonging to each taxa grouping in our study, we conducted cross-correlations among taxa to guide further study. Clusters of functionally-related taxa that exhibit correlated population dynamics could be directly influenced by specific climatic conditions or affected by the same biological feed-backs. In contrast, a negative correlation among taxa could indicate species interactions or opposing responses to external factors such as climate. Cross-correlation analysis across habitats revealed frequent significant correlations among taxa (Figure 3). Significant negative correlations (n=13) were almost as common as significant positive correlations (n=15). Collembola, parasitoids (Chalcidoidea) and Culicidae were typically negatively correlated to other taxa, while Collembola, Muscidae, and linyphiid spiders were positively correlated to each other. Lycosid spiders were positively correlated to other predators and parasitoids, but negatively to Collembola and Culicidae (Figure 3). Cross-correlation analyses within each individual habitat are shown in Figure S2.

**Figure 3.**
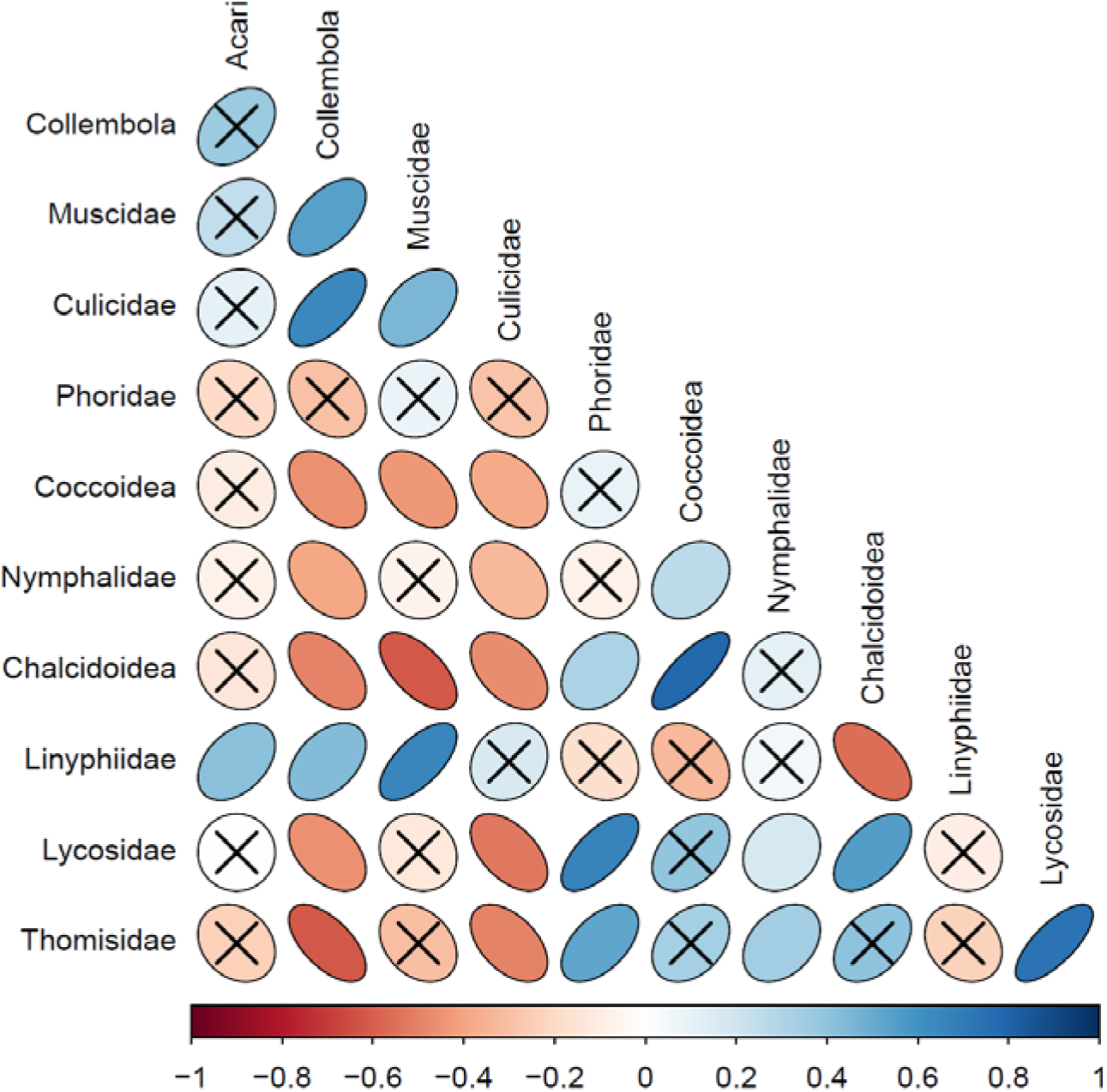
Cross-correlations of family or higher taxonomic level abundances across habitats based on non-parametric Spearman rank correlations. Ellipsoid shape indicates the strength of the spearman rank correlation, with more narrow ellipses depicting stronger cross-correlations between groups (red for negative and blue for positive correlations; the bar at the bottom of the graph depicts colour coding for the correlation coefficients). Crosses between groups depict those with non-significant correlations at the p < 0.05 level. The families are organized according to broad functional groups as outlined in Table S1. Five taxa which were not significantly correlated to any other groups are removed from the figure for clarity (Aphidoidea, Chironomidae, Ichneumonidae, Scatophagidae, and Sciaridae).

Finally, in order to gain insight into how trends for broader taxonomic groups relate to dynamics at the species level, we compared species-specific trends with their corresponding family-level trends. Specifically, using species-level data for all species in the spider family Linyphiidae and the fly family Muscidae, we estimated the temporal trends in abundance through time at both taxonomic levels (Figure 4). We found that there were more pronounced and contrasting abundance trends among the individual species of muscid flies as compared to the overall trend for the family (Figure S3). By contrast, the family-level trend for linyphiids largely resembled the trends of individual species (Figure S4). Generally, trends were most variable among species in the arid habitats (Figure 4).

**Figure 4.**
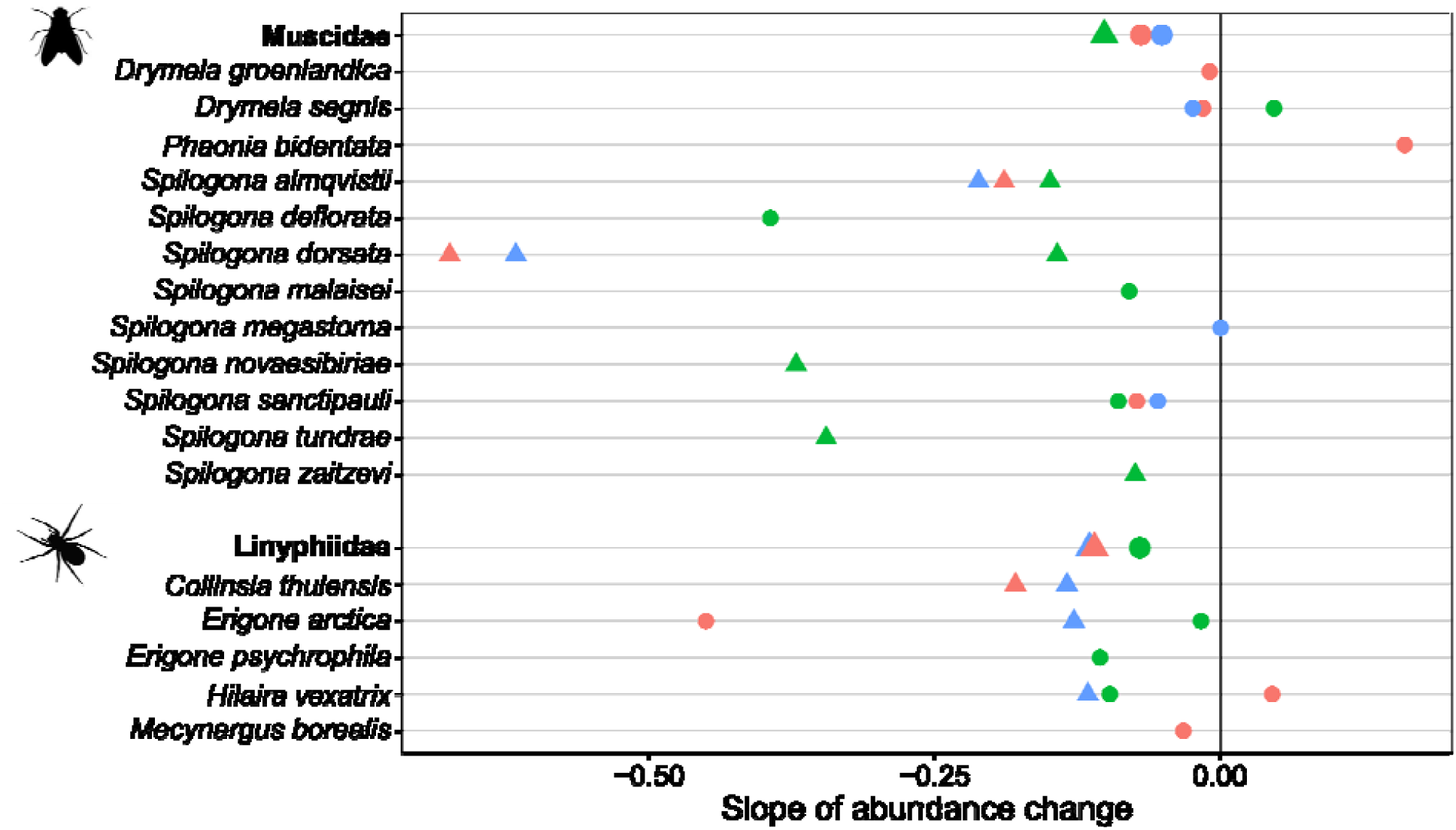
Species-level trends compared to family-level trends for the spider family Linyphiidae and the fly family Muscidae for which multiple species are represented in the dataset and species-level data was available. Temporal trends are estimated with the non-parametric modified Mann-Kendall trend analysis and significant trends at p < 0.05 are indicated by triangles. Symbol colours are the same as in Figure 1.

## DISCUSSION

Recent commentaries of insect declines have called for standardised methodology, well-designed syntheses of insect demography (31), balanced analysis (14), and realistic interpretation of results given the limitations of available data (32). In this study, we attempt to meet these challenges by presenting all relevant data from a long-term, standardised monitoring programme in the Arctic whereby arthropods were sampled weekly across the entire growing season. As a result of this effort, we identify substantial gaps in our knowledge of arthropod abundance trends. For example, based upon previous work from Zackenberg documenting effects of climate on body size (33, 34), phenology (35-37), abundance (16, 38), community composition (28, 30) and species interactions (39, 40), we expected there to be clear links between trends in climate and arthropod abundances, but we actually found much more complex dynamics, even at a broad taxonomic resolution of functional groups. While we did find some negative trends in arthropod abundance and even evidence that they are becoming more common, the abundances of most taxa show few directional changes and display substantial inter-annual variability. We note that climatic parameters also exhibit substantial inter-annual variability. Furthermore, our results from the cross-correlation analysis highlight the inherent challenges in attempting to understand complex interactions among taxa from long-term monitoring data. We also caution that species-level abundance trends may not be reflected in trends of broader taxonomic groups. Together, our findings suggest that we may not have successfully captured the relevant climate drivers, that species interactions and density dependent feedbacks could play stronger roles than have been previously acknowledged, that broad taxonomic groups are often comprised of species that do not fluctuate in synchrony or a combination of the above. Still, the empirical findings that we present here could have only emerged from long-term, standardized data, and we argue that it is by integrating such data with experiments and modelling that we will be best prepared to advance our mechanistic understanding of arthropod population dynamics (41).

### Long-term dynamics across taxa and habitats

The rapid climatic changes and the absence of many of the other human-mediated disturbances should make it easier to attribute long-term changes in Arctic arthropod population dynamics to shifts in environmental conditions, compared to temperate regions. However, it is also clear that arthropod population dynamics typically exhibit substantial idiosyncratic inter-annual variation, even in ecosystems primarily driven by climate (16, 28, 29). In addition, differences between short-term and long-term trends (42), as well as the occurrence of extreme events (43) are important to keep in mind as we develop a more synthetic understanding of the global phenomenon of insect declines (17).

Despite the range of temporal patterns we found here, the change point analysis provides insight into the nature of the trends. Change in local climate is not linear and the different climate factors do not change in synchrony. Most notably, in 2014 the change point for the climate PC1 (warmer falls and higher summer precipitation; Figure 2), coincides with significant change points for most taxa. The recent mix of positive and negative changes are congruent with the trends found in the most recent short term time series (Figure 1d), and thereby suggests that there are winners and losers under certain changing climate conditions. However, many of the earlier change points in arthropod abundance are more difficult to summarise, because they do not coincide with climate changes, indicating that long-term trends can be complicated by inter-annual and multi-annual variability. We note that individual species within taxa could be responding differently, and that although rare taxa were omitted, the likelihood of detecting change points is possibly lower for the less abundant taxa included in our analyses.

A number of challenges remain in linking temporal variation in environmental conditions to arthropod populations. The first and arguably most important is a better understanding of which climate metrics are the most relevant for predicting responses to change in individual arthropod species and even life stages. Here, the choice of location for temperature measurements (air, surface, or soil), the spatial resolution, and the choice of metric (mean, extreme, or variability) need to be further explored. Water stress is another known factor of importance to arthropods in cold environments (44), which may interact with temperature stress (45). Although soil moisture varies over small spatial scales (46), and thus can be challenging to measure, such variation has been shown to influence arthropod communities (47, 48). We are also only beginning to understand how thermal tolerance of Arctic arthropods varies across life stages (e.g. 49) and in response to past exposure to extreme conditions (e.g. 50). Variation in climate-abundance relationships may be further affected by the predictability of the thermal and hydrological environment (51), and flexible life stage lengths and generation times could buffer unfavourable conditions. Such complexities are likely to be even greater further south, where anthropogenic pressures are more pronounced. New efforts to compile and model microclimate data may help in our understanding of relevant scales (52, 53), but much more data from cold biomes are highly needed and called for (e.g., by the SoilTemp initiative 54).

### Climate effects vs. biotic feed-back mechanisms

Strong relationships with climate drivers have been identified for life history variation in individual species in the past (33, 34, 36). However, the results from our cross-correlation analyses clearly demonstrate that many taxa do not fluctuate in synchrony or even exhibit strong negatively correlated dynamics. For example, the negative correlation and the opposing change point patterns between Lycosidae and Collembola could arise from their trophic link to each other, but indirect climate impacts may also independently impact these groups. For example, Collembola abundances may decline as has been found in warming experiments (55), with concurrent increases in Lycosidae due to activity-related improved hunting ability (56) or via more complex indirect interactions (57, 58). Similarly, positive correlations among taxa with related diets (Lycosidae and Thomsidae) could indicate co-existence via niche diversity, mutualisms or independent responses to biotic or abiotic drivers (59). The cause of negative correlations with parasitoids and Culicidae remains unclear but could be related to their distinct life cycles which do not resemble those of the majority of other taxa. We also note that correlations or lack thereof, may mask substantial variation in correlations among individual species. In any case, taxa that are correlated in abundance provide interesting focal points for future work. To advance the field of insect population dynamics, we need to support more basic work on the resource requirements, climatic constraints, and biological feed-backs of tundra arthropod species and their interactions (15, 40, 60).

### Abundance changes at the species-level

Trait-based approaches allow for broader functional inference about species-environment relationships (55, 61). However, such approaches rely on accurate and complete information about functionally important traits, which is severely lacking for most arthropods (62). For practical reasons, species are often lumped together in broad taxonomic groups, which may not be functionally similar. In some cases, higher taxonomic resolution can be sufficient to understand overall trends in community structure, or biomass (30, 63), but our study suggests that this is often not the case for variation in abundance. There is clearly also a problem with feasibility in running long-term arthropod monitoring programs and resolving time series to the species level, which explains why there are very few long-term arthropod monitoring programs worldwide. However, our study demonstrates that when we aggregate species to broader taxonomic levels, our ability to understand the drivers of abundance change are constrained, as even closely related species may differ in their response to environmental drivers (28, 29).

### Recommendations for future work

Arthropod monitoring in remote regions such as the Arctic is rare, yet critical to our understanding of the role of climate in arthropod population dynamics. New monitoring initiatives should align with established programs to ensure inter-site comparability with respect to sampling methodology, spatio-temporal coverage, curation of samples and specimens, and digitization of collections (15, 31, 64). More efforts should also be made to address the common problems with monitoring and analysing arthropod trends described at length by Didham, *et al.* (32). Complementary to this, large-scale programs utilizing standardized sampling protocols and covering vast gradients in environmental conditions should be implemented (15). However, both approaches are hampered by our current lack of sufficient taxonomic resolution, our limited knowledge about which life-history stages, populations, and species are the most sensitive to environmental change, as well as how species interact. One solution to this is to focus on a few taxa that may serve as indicators. However, given that species respond differently to environmental drivers (15, 28), we need better data on species-specific trends before selecting potential indicator species. Arthropod trait databases could contribute to a better understanding of which species are likely to exhibit correlated responses to climatic factors (61, 62). We acknowledge that it is often not feasible to obtain all this information due to lack of expertise and funding. However, current technological developments offer promising outlooks. For instance, molecular tools can provide high taxonomic resolution, and although abundance thought to be difficult to derive, new analytical tools are emerging (e.g. 65). Image-based solutions employing deep learning models form a powerful tool for automated insect identification and biomass estimation as well as for monitoring of species and species interactions in their natural environment (66, 67). In addition to improved taxonomic resolution and trait information, a current and major challenge is that there is limited data collection of relevant environmental conditions that are likely driving arthropod abundance changes. Such data should be collected at the localized scale of arthropod sampling, including at established monitoring stations where such data have thus far been missing. From there, relevant modelling tools can be further developed to scale-up paired arthropod and micro-climatic data. Together such future advances to our understanding of arthropod population dynamics require interdisciplinary efforts such as those stimulated by the Network for Arthropods of the Tundra (60).

### Conclusions

This study is the product of a relatively simple monitoring program in the High Arctic, and demonstrates that data from such a site can provide invaluable insights into arthropod population trends when collected over long time periods. We find that even in a species-poor ecosystem subject to rapid climate change but few other direct human disturbances, arthropod population dynamics are no less challenging to comprehend. Unlike recent high profile findings further south, we have not found evidence of an impending “insectageddon” (*sensu* 14), although some taxa and individual species exhibit strong and increasing negative trends in abundance. Nevertheless, the complexities we have shown highlight the need for detailed long-term data and thoughtful analysis. For example, we require even longer term data to establish whether our change point patterns are random, are biological records of extreme events, or whether they represent longer cyclic patterns in abundance (e.g., via density dependent mechanisms or interspecific interactions). We need more such insights from a wider range of locations around the Arctic region, which could form the basis of comparable datasets. We suggest that biome-wide coordination of monitoring efforts, species-level population and trait data improvements, the continual development of efficient monitoring technologies, and the collection of relevant climate variables are equally important in the pursuit of a fuller understanding of biotic and abiotic impacts on such a pivotal group of organisms.

## MATERIALS AND METHODS

### Sampling and specimens

The Greenland Ecosystem Monitoring Programme has been in operation at Zackenberg in Northeast Greenland (74°28’N, 20°34’W) since 1996, and includes standardized pitfall trapping in five plots: one fen, two mesic heaths, and two arid heath plots (see 23 for further details). Each plot consisted of eight yellow pitfall traps (1997 – 2006) or four pitfall traps (1996 & 2007 – 2018; each trap 5m from nearest neighbour). The traps were in operation during the growing season starting at snowmelt in late May-early June and ending by late August-early September, and emptied once a week. To compare across the years of sampling (1996–2018), we truncated our arthropod data to June, July, and August. Arthropods were then sorted to the family level for spiders and most insects, superfamily level for Aphidoidea, Chalcidoidea, and Coccoidea, and subclass level for other arthropods and counted as part of the monitoring programme. We note that juveniles were included in the family level abundances for spiders.

For the years 1996–2014, all spiders, as well as the muscid flies from selected traps, were identified to the species. For species-level data, only adults were used in analyses as juvenile spiders and fly larvae could have different dynamics, and are difficult to identify with confidence. For spiders, we included individuals sampled in all five plots, whereas for muscid flies, only a subset of traps from the wet fen plot, one mesic and one arid heath plot were identified (see 28 for details). For some early years of the programme, certain families of Diptera were not sorted, but one family strongly dominated samples from later years (35), hence we pooled the Chironomidae and Ceratopogonidae (hereafter called Chironomidae), Anthomyiidae and Muscidae (hereafter called Muscidae) and Mycetophilidae and Sciaridae (hereafter called Sciaridae), respectively.

The wet fen plot is primarily composed of mosses, grasses (e.g., *Eriophorum* sp.), and sedges with scattered Arctic willow (*Salix arctica*). The mesic plots primarily consist of Arctic bell-heather (*Cassiope tetragona*) and Arctic willow, grasses/sedges, and berry plants (*Vaccinium*), while the arid plots have a greater dominance of mountain avens (*Dryas octopetala*) (35). All plots are located within an area less than one km^2^. The wet fen and arid plots typically have snowmelt two weeks earlier than the mesic plots and soil moisture is highest in the wet fen and lowest in the arid plots. Air temperature (2 m above the surface), soil temperature (at 0cm, 5cm and 10cm depth) in mesic heath dominated by *Cassiope tetragona*, and precipitation (mm) was measured hourly by a meteorological station located centrally to all the arthropod sampling plots (68).

### Data and analyses

We standardized weekly abundance counts for each arthropod group by calculating the abundance per trap per day for each plot in each season. In subsequent analyses, we focused on taxonomic groups for which total abundance in each habitat across the time series was higher than 500 individuals to ensure robust estimates of trends even for taxa that do not occur in all years. We derived a range of climate variables of relevance to arthropod populations from the temperature and precipitation data. Summer precipitation was calculated from the sum of June, July, and August measurements. Average summer temperature was calculated from air temperature measured during the same period (June, July, and August). Most Arctic arthropods overwinter beneath the snow in leaf litter and the upper portion of the soil profile. Thus, in order to estimate average seasonal temperatures experienced by arthropods for the previous fall (October_t-1_ and November_t-1_), winter (December_t-1_ through March_t_), and spring (April_t_ and May_t_), we calculated the mean hourly temperature across the data collected at 0 cm, 5 cm, and 10 cm depth within the soil profile. We defined the length of winter as the number of days from the first day in the fall with an average temperature below 2°C to the last day in spring with an average temperature below 2°C. The number of freeze-thaw cycles over the previous winter were calculated from the soil temperature data by counting the number of days with a maximum temperature above 0°C and a minimum temperature below 0°C. To avoid statistical artefacts associated with multicollinearity (69), we reduced the various climatic variables in our time series to a smaller number of composite predictors by doing a principal components analysis (PCA) using the factoMineR package (70). In our PCA, we also included lagged variables for summer precipitation (precipitation_t-1_) and summer temperature (summer_t-1_), as conditions during the previous growing season could influence arthropod abundance in summer_t_. Precipitation data were log-transformed and all climate variables were centred and scaled prior to PCA.

Temporal trends in abundance were analyzed by taxa and habitat with the Mann-Kendall trend test corrected for temporal autocorrelation (71) using the *fume* package for the corrected Mann-Kendall trend test (72). We applied a change-point detection approach to each time series (climate and arthropod abundance) to identify potential shifts over time. We used the ‘breakpoints’ function of the *strucchange* package (73) to test the null hypothesis that no abrupt change has occurred over time. We compared several simple regression models with no change and with different numbers of change-points across years. The F-statistic procedure identifies the best regression model with the smallest residual sum of squares based on the Bayesian Information Criterion (BIC) (74). Cross correlation in abundance among the taxonomic groups were estimated by non-parametric spearman rank correlations and visualized using the *corrplot* package (75). All statistical analyses were performed in the R 3.6.1 platform (76).

## ACKNOWLEDGEMENTS

David Wagner is gratefully thanked for convening the session “Insect declines in the Anthropocene” at the Entomological Society of America annual meeting 2019 in St. Louis, USA, which brought the group of contributors to the special feature together. TTH acknowledges funding from the Independent Research Fund Denmark (grant 8021-00423B). Data were kindly provided by the Greenland Ecosystem Monitoring programme. We thank the Danish Environmental Protection Agency for funding over the years.

## REFERENCES

1. J. C. Habel et al., Butterfly community shifts over two centuries. Conserv. Biol. 30, 754–762 (2016).

2. S. Seibold et al., Arthropod decline in grasslands and forests is associated with landscape-level drivers. Nature 574, 671–674 (2019).

3. C. A. Hallmann et al., More than 75 percent decline over 27 years in total flying insect biomass in protected areas. PLOS ONE 12, e0185809 (2017).

4. J. C. Biesmeijer et al., Parallel declines in pollinators and insect-pollinated plants in Britain and the Netherlands. Science 313, 351–354 (2006).

5. L. G. Carvalheiro et al., Species richness declines and biotic homogenisation have slowed down for NW-European pollinators and plants. Ecol. Lett. 16, 870–878 (2013).

6. D. Maes, H. Van Dyck, Butterfly diversity loss in Flanders (north Belgium): Europe’s worst case scenario? Biol. Conserv. 99, 263–276 (2001).

7. J. A. Thomas et al., Comparative losses of British butterflies, birds, and plants and the global extinction crisis. Science 303, 1879–1881 (2004).

8. C. R. Shortall et al., Long-term changes in the abundance of flying insects. Insect. Conserv. Diver. 2, 251–260 (2009).

9. R. Dirzo et al., Defaunation in the Anthropocene. Science 345, 401–406 (2014).

10. B. C. Lister, A. Garcia, Climate-driven declines in arthropod abundance restructure a rainforest food web. Proceedings of the National Academy of Sciences 115, E10397–E10406 (2018).

11. W. E. Kunin, Robust evidence of insect declines. Nature 574, 641–642 (2019).

12. A. Komonen, P. Halme, J. S. Kotiaho, Alarmist by bad design: Strongly popularized unsubstantiated claims undermine credibility of conservation science. Rethinking Ecology 4 (2019).

13. M. R. Willig et al., Populations are not declining and food webs are not collapsing at the Luquillo Experimental Forest. Proc. Natl. Acad. Sci. USA 116, 12143–12144 (2019).

14. C. D. Thomas, T. H. Jones, S. E. Hartley, “Insectageddon”: A call for more robust data and rigorous analyses. Glob. Change Biol. 25, 1891–1892 (2019).

15. M. A. K. Gillespie et al., Circumpolar terrestrial arthropod monitoring: A review of ongoing activities, opportunities and challenges, with a focus on spiders. Ambio 49, 704–717 (2020).

16. M. A. K. Gillespie et al., Status and trends of terrestrial arthropod abundance and diversity in the North Atlantic region of the Arctic. Ambio 49, 718–731 (2020).

17. D. L. Wagner, Insect declines in the Anthropocene. Annu. Rev. Entomol. 65, 457–480 (2020).

18. F. Sánchez-Bayo, K. A. G. Wyckhuys, Worldwide decline of the entomofauna: A review of its drivers. Biol. Conserv. 232, 8–27 (2019).

19. P. Soroye, T. Newbold, J. Kerr, Climate change contributes to widespread declines among bumble bees across continents. Science 367, 685–688 (2020).

20. M. A. K. Gillespie et al., A method for the objective selection of landscape-scale study regions and sites at the national level. Methods Ecol Evol 8, 1468–1476 (2017).

21. ABA, Arctic Biodiversity Assessment - status and trends of arctic biodiversity (Conservation of Arctic Flora and Fauna, Arctic Council, 2013).

22. IPCC, Climate Change 2014: Impacts, Adaptation, and Vulnerability. Part B: Regional Aspects. Contribution of Working Group II to the Fifth Assessment Report of the Intergovernmental Panel on Climate Change (Cambridge University Press, Cambridge, United Kingdom and New York, NY, USA, 2014), pp. 688.

23. N. M. Schmidt, L. H. Hansen, J. Hansen, T. B. Berg, H. Meltofte, BioBasis - Conceptual design and sampling procedures of the biological monitoring programme within Zackenberg Basic, pp. 107, (2019).

24. T. Brereton, D. B. Roy, I. Middlebrook, M. Botham, M. Warren, The development of butterfly indicators in the United Kingdom and assessments in 2010. J. Insect Conserv. 15, 139–151 (2011).

25. Y. Melero, C. Stefanescu, J. Pino, General declines in Mediterranean butterflies over the last two decades are modulated by species traits. Biol. Conserv. 201, 336–342 (2016).

26. D. R. Brooks et al., Large carabid beetle declines in a United Kingdom monitoring network increases evidence for a widespread loss in insect biodiversity. J. Appl. Ecol. 49, 1009–1019 (2012).

27. H. Van Dyck, A. J. Van Strien, D. Maes, C. A. M. Van Swaay, Declines in common, widespread butterflies in a landscape under intense human use. Conserv. Biol. 23, 957–965 (2009).

28. S. Loboda, J. Savage, C. M. Buddle, N. M. Schmidt, T. T. Høye, Declining diversity and abundance of High Arctic fly assemblages over two decades of rapid climate warming. Ecography 41, 265–277 (2018).

29. J. J. Bowden, O. L. P. Hansen, K. Olsen, N. M. Schmidt, T. T. Hoye, Drivers of inter-annual variation and long-term change in High-Arctic spider species abundances. Polar Biol. 41, 1635–1649 (2018).

30. A. M. Koltz, N. M. Schmidt, T. T. Høye, Differential arthropod responses to warming are altering the structure of Arctic communities. Royal Society Open Science 5 (2018).

31. G. A. Montgomery et al., Is the insect apocalypse upon us? How to find out. Biol. Conserv. 241, 108327 (2020).

32. R. K. Didham et al., Interpreting insect declines: seven challenges and a way forward. Insect. Conserv. Diver. 13, 103–114 (2020).

33. T. T. Høye, J. U. Hammel, T. Fuchs, S. Toft, Climate change and sexual size dimorphism in an arctic spider. Biol. Lett. 5, 542–544 (2009).

34. J. J. Bowden et al., High-Arctic butterflies become smaller with rising temperatures. Biol. Lett. 11, 20150574 (2015).

35. T. T. Høye, M. C. Forchhammer, Phenology of high-arctic arthropods: effects of climate on spatial, seasonal and inter-annual variation. Adv. Ecol. Res. 40, 299–324 (2008).

36. T. T. Høye et al., Phenology of high-arctic butterflies and their floral resources: Species-specific responses to climate change. Curr. Zool. 60, 243–251 (2014).

37. T. T. Høye, E. Post, H. Meltofte, N. M. Schmidt, M. C. Forchhammer, Rapid advancement of spring in the High Arctic. Curr. Biol. 17, R449–R451 (2007).

38. T. T. Høye, E. Post, N. M. Schmidt, K. Trøjelsgaard, M. C. Forchhammer, Shorter flowering seasons and declining abundance of flower visitors in a warmer Arctic. Nature Clim. Change 3, 759–763 (2013).

39. N. M. Schmidt et al., An ecological function in crisis? The temporal overlap between plant flowering and pollinator function shrinks as the Arctic warms. Ecography 39, 1250–1252 (2016).

40. N. M. Schmidt et al., Interaction webs in arctic ecosystems: Determinants of arctic change? Ambio 46, 12–25 (2017).

41. N. M. Schmidt, T. R. Christensen, T. Roslin, A high arctic experience of uniting research and monitoring. Earth’s Future 5, 650–654 (2017).

42. A. M. Iler, T. T. Hoye, D. W. Inouye, N. M. Schmidt, Long-Term Trends Mask Variation in the Direction and Magnitude of Short-Term Phenological Shifts. Am. J. Bot. 100, 1398–1406 (2013).

43. N. M. Schmidt, J. Reneerkens, J. H. Christensen, M. Olesen, T. Roslin, An ecosystem-wide reproductive failure with more snow in the Arctic. PLoS Biol. 17, e3000392 (2019).

44. P. Convey, W. Block, H. J. Peat, Soil arthropods as indicators of water stress in Antarctic terrestrial habitats? Glob. Change Biol. 9, 1718–1730 (2003).

45. M. J. Everatt, P. Convey, J. S. Bale, M. R. Worland, S. A. L. Hayward, Responses of invertebrates to temperature and water stress: A polar perspective. J. Therm. Biol. 54, 118–132 (2015).

46. P. C. le Roux, J. Aalto, M. Luoto, Soil moisture’s underestimated role in climate change impact modelling in low-energy systems. Glob. Change Biol. 19, 2965–2975 (2013).

47. R. R. Hansen et al., Meter scale variation in shrub dominance and soil moisture structure Arctic arthropod communities. Peerj 4, ARTN e2224 (2016).

48. T. T. Høye et al., Elevation modulates how Arctic arthropod communities are structured along local environmental gradients. Polar Biol. 41, 1555–1565 (2018).

49. S. E. Anthony, C. M. Buddle, T. T. Høye, B. J. Sinclair, Thermal limits of summer-collected Pardosa wolf spiders (Araneae: Lycosidae) from the Yukon Territory (Canada) and Greenland. Polar Biol. 42, 2055–2064 (2019).

50. M. H. Sørensen et al., Rapid induction of the heat hardening response in an Arctic insect. Biol. Lett. 15 (2019).

51. J. A. Deere, B. J. Sinclair, D. J. Marshall, S. L. Chown, Phenotypic plasticity of thermal tolerances in five oribatid mite species from sub-Antarctic Marion Island. J. Insect Physiol. 52, 693–700 (2006).

52. I. Bramer et al., Advances in Monitoring and Modelling Climate at Ecologically Relevant Scales. Next Generation Biomonitoring, Pt 1 58, 101–161 (2018).

53. M. R. Kearney, P. K. Gillingham, I. Bramer, J. P. Duffy, I. M. D. Maclean, A method for computing hourly, historical, terrain-corrected microclimate anywhere on earth. Methods Ecol Evol 11, 38–43 (2020).

54. J. J. Lembrechts, Etal., SoilTemp: call for data for a global database of near-surface temperature. Glob. Change Biol. (in press).

55. M. Makkonen et al., Traits explain the responses of a sub-arctic Collembola community to climate manipulation. Soil Biol. Biochem. 43, 377–384 (2011).

56. A. L. Asmus et al., Shrub shading moderates the effects of weather on arthropod activity in arctic tundra. Ecol. Entomol. 43, 647–655 (2018).

57. A. M. Koltz, A. T. Classen, J. P. Wright, Warming reverses top-down effects of predators on belowground ecosystem function in Arctic tundra. Proceedings of the National Academy of Sciences 115, E7541–E7549 (2018).

58. A. M. Koltz, J. P. Wright, Impacts of female body size on cannibalism and juvenile abundance in a dominant arctic spider. J. Anim. Ecol. 10.1111/1365-2656.13230 (in press).

59. H. K. Wirta, E. Weingartner, P. A. Hamback, T. Roslin, Extensive niche overlap among the dominant arthropod predators of the High Arctic. Basic Appl. Ecol. 16, 86–92 (2015).

60. T. T. Høye, L. E. Culler, Tundra arthropods provide key insights into ecological responses to environmental change. Polar Biol. 41, 1523–1529 (2018).

61. M. K. L. Wong, B. Guénard, O. T. Lewis, Trait-based ecology of terrestrial arthropods. Biological Reviews 94, 999–1022 (2019).

62. M. M. Gossner et al., A summary of eight traits of Coleoptera, Hemiptera, Orthoptera and Araneae, occurring in grasslands in Germany. Scientific Data 2, 150013 (2015).

63. L. L. Timms, J. J. Bowden, K. S. Summerville, C. M. Buddle, Does species-level resolution matter? Taxonomic sufficiency in terrestrial arthropod biodiversity studies. Insect. Conserv. Diver. 6, 453–462 (2013).

64. D. S. Sikes et al., The value of museums in the production, sharing, and use of entomological data to document hyperdiversity of the changing North. Arctic Science 3, 498–514 (2017).

65. Y. Ji et al., SPIKEPIPE: A metagenomic pipeline for the accurate quantification of eukaryotic species occurrences and intraspecific abundance change using DNA barcodes or mitogenomes. Mol Ecol Resour 20, 256–267 (2020).

66. J. Ärje et al., Automatic image-based identication and biomass estimation of invertebrates. ArXiv, 2002.03807 (2020).

67. K. Bjerge, M. V. Sepstrup, J. B. Nielsen, F. Helsing, T. T. Hoye, A light trap and computer vision system to detect and classify live moths (Lepidoptera) using tracking and deep learning. bioRxiv 10.1101/2020.03.18.996447, 2020.2003.2018.996447 (2020).

68. N. Kandrup, K. M. Iversen, Zackenberg Ecological Research Operation - ClimateBasis Manual, pp. 15, Project no: B15 (2010).

69. C. F. Dormann et al., Collinearity: a review of methods to deal with it and a simulation study evaluating their performance. Ecography 36, 27–46 (2013).

70. S. Lê, J. Josse, F. Husson, FactoMineR: An R Package for Multivariate Analysis. 2008 25, 18 (2008).

71. G. Patle, A. Libang, S. Ahuja (2016) Analysis of rainfall and temperature variability and trend detection: A non parametric Mann Kendall test approach. in 2016 3rd International Conference on Computing for Sustainable Global Development (INDIACom) (IEEE), pp 1723–1727.

72. S. M. Group, FUME package, pp. (2012).

73. A. Zeileis, F. Leisch, K. Hornik, C. Kleiber, strucchange. An R package for testing for structural change in linear regression models. Journal of Statistical Software 7, 1–38 (2002).

74. A. Zeileis, C. Kleiber, W. Krämer, K. Hornik, Testing and dating of structural changes in practice. Comput. Stat. Data Anal. 44, 109–123 (2003).

75. T. Wei, V. Simko (2017) R package ‘corrplot’: Visualization of a correlation matrix.

76. R Core Team (2019) R: A langage and environment for statistical computing. (R Foundation for Statistical Computing, Vienna, Austria).

