## Supplementary information for "Taxonomic, temporal, and spatial variation in the dynamics of High-Arctic arthropod populations"

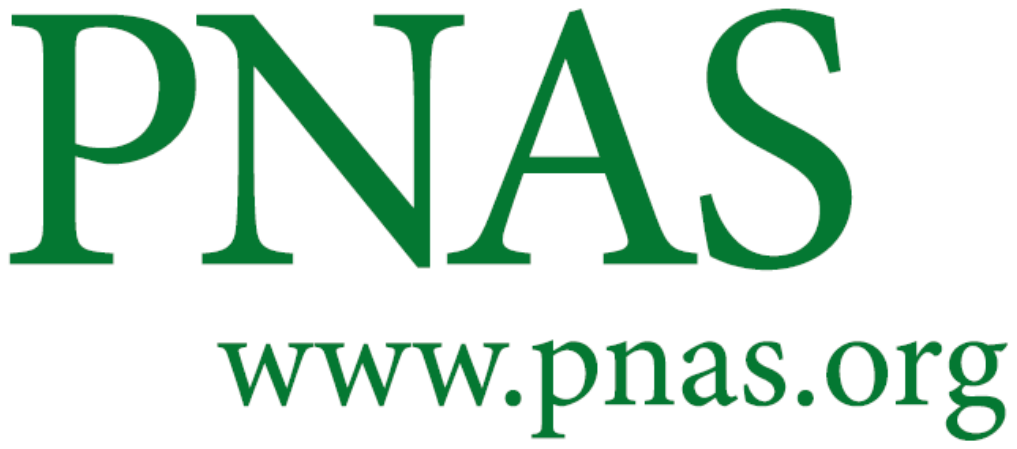

Supplementary Information for

Taxonomic, temporal, and spatial variation in the dynamics of High-Arctic arthropod populations

Toke T. Høye, Sarah Loboda, Amanda M. Koltz, Mark A. K. Gillespie, Joseph J. Bowden, Niels M. Schmidt

Toke T. Høye

**This PDF file includes:**

Figures S1 to S4

Tables S1 to S5

Figure S1

Annual variation in climate variables included in the Principal Component Analysis to describe local climatic variation at Zackenberg, Greenland, between 1996 and 2018. Regression fits are based on change point analysis. Detailed results of change point analysis are presented in Table S3.

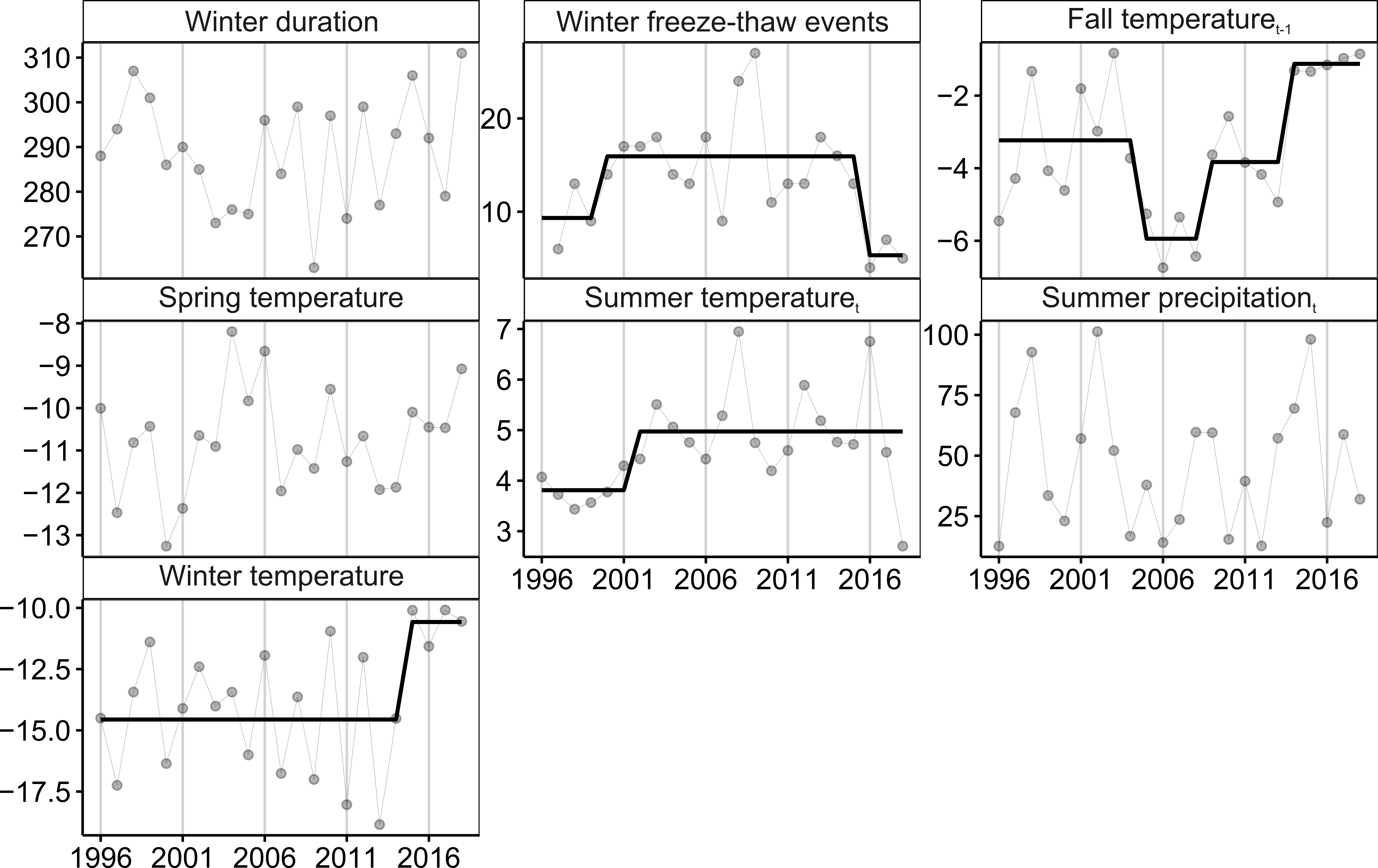

Figure S2

Cross-correlations of family- or higher taxa level abundances in each habitat based on non-parametric Spearman rank correlations. Ellipsoid shape indicates the strength of the spearman rank correlation, with more narrow ellipses depicting stronger cross-correlations between groups (red for negative and blue for positive correlations; the bar at the bottom of the graph depicts colour coding for the correlation coefficients). Crosses between groups depict those with non-significant correlations at the p < 0.05 level. Taxa for which no cross-correlations were found are omitted for clarity.

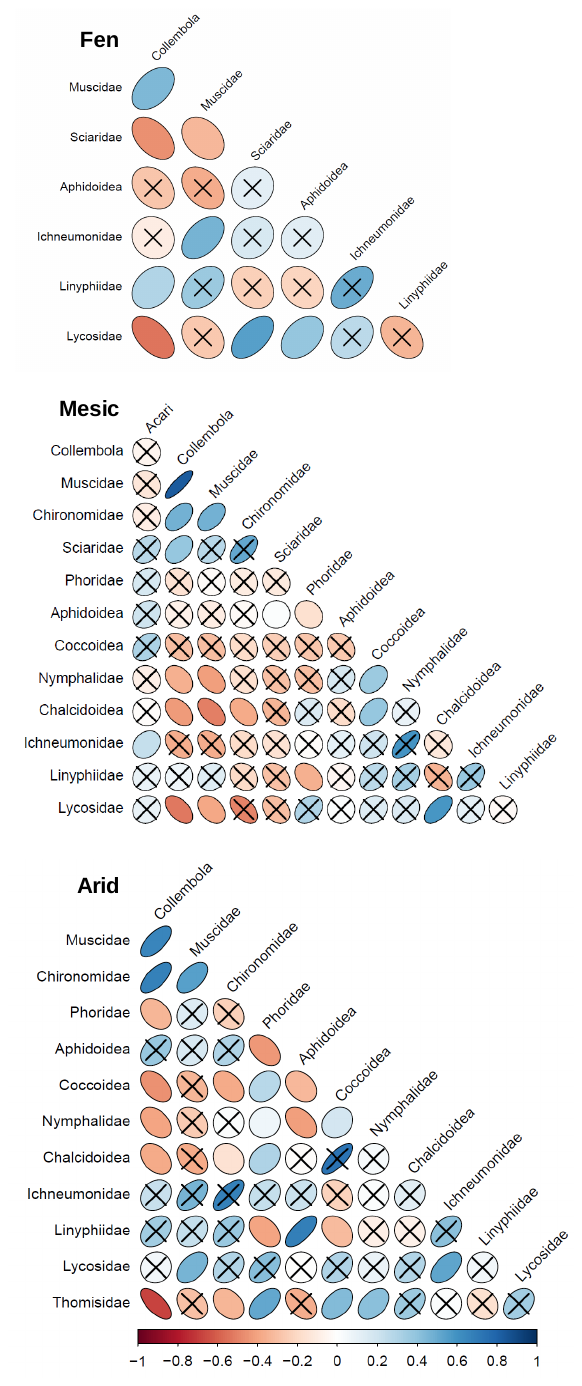

Figure S3

Time series of annual abundances and trends over time for each species of fly in the family Muscidae; the last panel shows the total abundance per year and trend for the whole family (i.e., from combining data from the individual species). The time series are presented separately for each of the three habitat types (fen, mesic heath, arid heath).

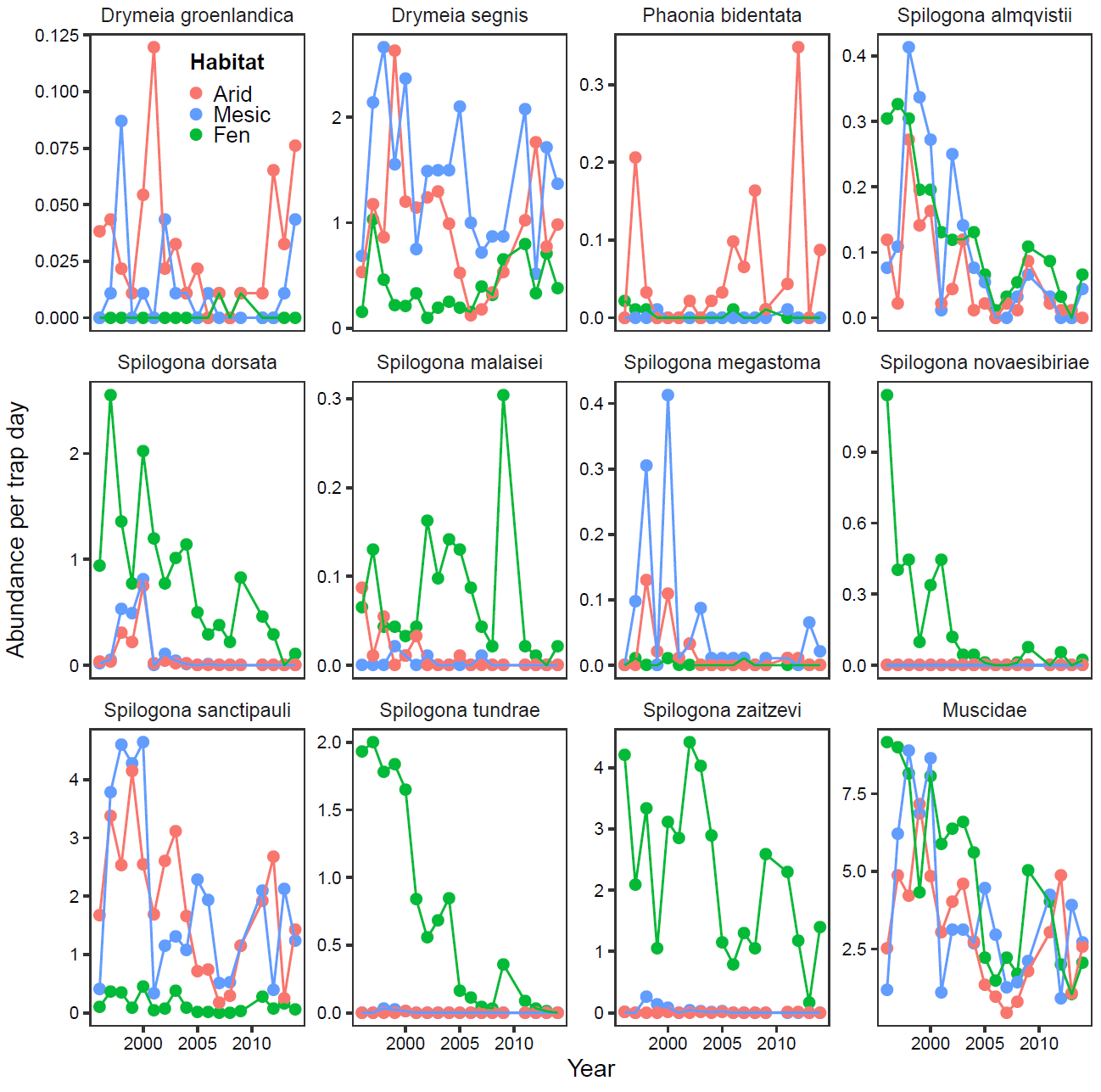

Figure S4

Time series of annual abundances and trends over time for each spider species in the family Linyphiidae; the last panel shows the total abundance per year and trend for the whole family (i.e., from combining data from the individual species). The time series are presented separately for each of the three habitat types (fen, mesic heath, arid heath).

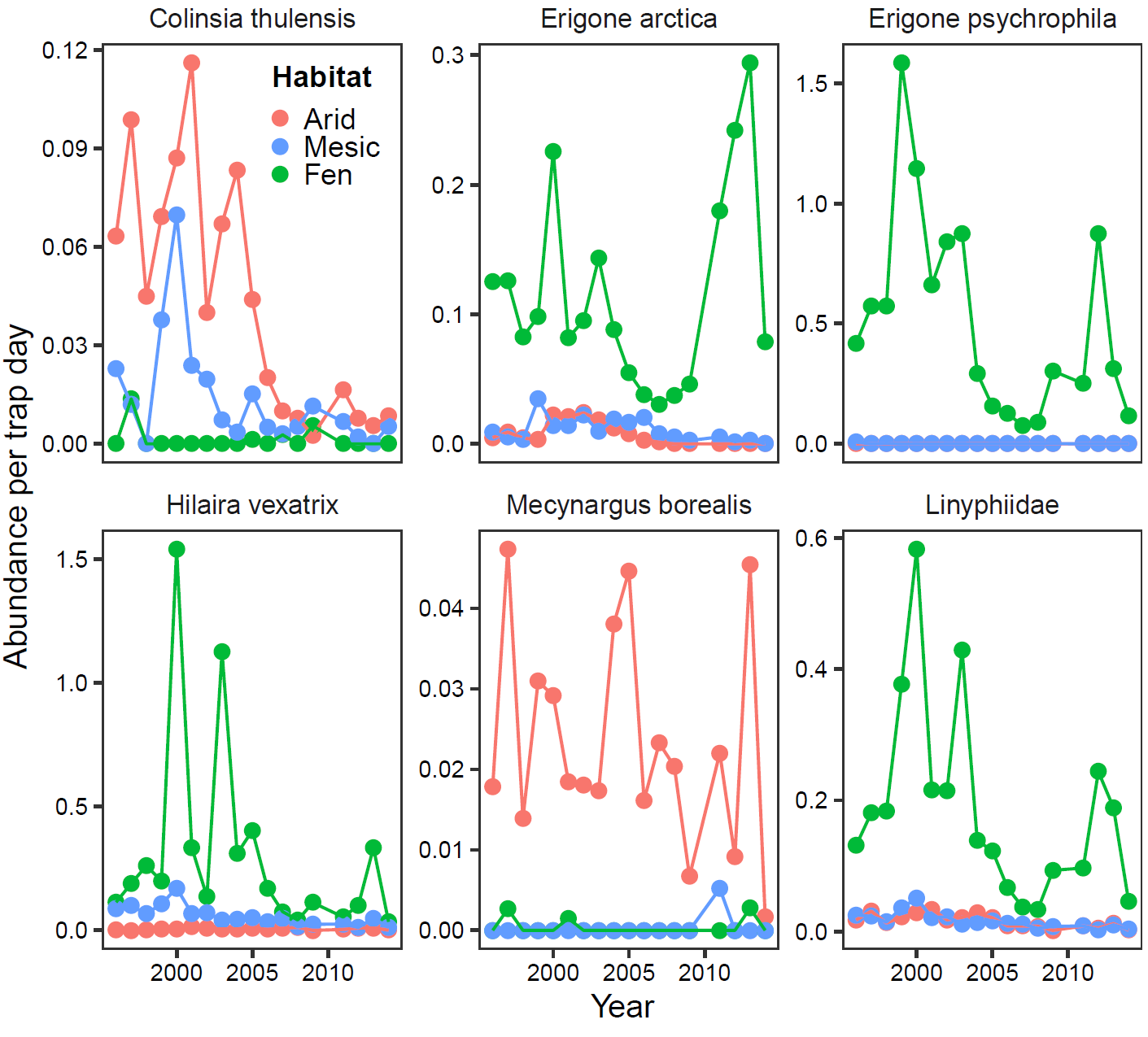

Table S1

Overview of the taxa included in this study, their taxonomic affiliations, and our assignment to broad functional groups.

| **Order/subclass** | **Family/superfamily** | **Functional group** |
| --- | --- | --- |
| Acari |  | Detritivores |
| Collembola |  | Detritivores |
| Aranea | Linyphiidae | Predators |
| Aranea | Lycosidae | Predators |
| Aranea | Thomisidae | Predators |
| Diptera | Chironomidae | Mixed feeders |
| Diptera | Muscidae | Mixed feeders |
| Diptera | Phoridae | Mixed feeders |
| Diptera | Sciaridae | Mixed feeders |
| Diptera | Scatophagidae | Mixed feeders |
| Hemiptera | Aphidoidea | Herbivores |
| Hemiptera | Coccoidea | Herbivores |
| Hymenoptera | Chalcidoidea | Parasitoids |
| Hymenoptera | Ichneumonidae | Parasitoids |
| Lepidoptera | Nymphalidae | Pollinators |

Table S2

Loadings of individual climate variables from the Principal Components Analysis after varimax rotation, as well as the variation explained by each individual axis and the cumulative variation. Both precipitation variables were log transformed and all variables were scaled and centered prior to analysis.

| **Parameter** | **PC1** | **PC2** | **PC3** |
| --- | --- | --- | --- |
| Average fall temperature_t-1_ | 0.92 | 0.07 | 0.01 |
| Summer precipitation_t-1_ | 0.76 | -0.2 | 0.28 |
| Summer precipitation_t_ | 0.53 | 0.09 | -0.42 |
| Average summer temperature_t-1_ | 0.09 | 0.85 | 0.22 |
| Winter duration | 0.2 | -0.8 | 0.34 |
| Average spring temperature_t_ | -0.02 | 0.09 | 0.94 |
| Average winter temperature | 0.48 | -0.28 | 0.75 |
| Average summer temperature_t_ | 0.00 | 0.22 | -0.03 |
| Number of freeze-thaw events | -0.2 | 0.22 | -0.09 |
| Eigenvalue | 2.82 | 1.77 | 1.63 |
| Percentage of variance explained | 31.34 | 19.68 | 18.1 |
| Cumulative percentage of variance | 31.34 | 51.02 | 69.13 |

Table S3

Results of change point analysis for each of the raw climatic variables used in the PCA to describe environmental change at Zackenberg between 1996 and 2018 and for the three varimax-rotated PCs as well (Table S2 describes the loadings of the climatic variables onto each of the PCs). RSS= Residual sum of squares, BIC= Bayesian Information Criterion. RSS and BIC values for the model with the lowest BIC value are boldfaced.

|  |  | **Number of break dates in the model** | | | | | | | **Corresponding break dates** |
| --- | --- | --- | --- | --- | --- | --- | --- | --- | --- |
| **Variable** |  | **0** | **1** | **2** | **3** | **4** | **5** | **6** |  |
| Average spring temperature_t_ | RSS | **34.03** | 28.77 | 20.65 | 16.92 | 15.43 | 15.02 | 14.92 |  |
|  | BIC | **80.55** | 82.96 | 81.61 | 83.3 | 87.44 | 93.09 | 99.21 |  |
| Average winter temperature | RSS | 152.62 | **100.23** | 91.72 | 89.01 | 86.94 | 85.64 | 84.97 | 2014 |
|  | BIC | 115.07 | **111.67** | 115.9 | 121.48 | 127.21 | 133.13 | 139.23 |  |
| Average fall temperature_t-1_ | RSS | 78.18 | 45.8 | 35.17 | **25.24** | 19.45 | 22.85 | 28.07 | 2004, 2008, 2013 |
|  | BIC | 99.68 | 93.66 | 93.85 | **92.49** | 92.77 | 102.74 | 113.75 |  |
| Summer precipitation_t_ | RSS | **16894.6** | 15888.7 | 13580.3 | 11504.6 | 10404.1 | 9950.6 | 9165.5 |  |
|  | BIC | **223.3** | 228.2 | 230.8 | 233.3 | 237.3 | 242.5 | 246.9 |  |
| Freeze-thaw events | RSS | 748.4 | 531.6 | **354.6** | 318.6 | 271.5 | 253.5 | 262 | 1999, 2015 |
|  | BIC | 151.6 | 150 | **147** | 150.8 | 153.4 | 158.1 | 165.1 |  |
| Winter duration | RSS | **3421.8** | 3064.8 | 2503.5 | 2368.3 | 2117.5 | 1928.5 | 1836.3 |  |
|  | BIC | **186.6** | 190.3 | 192 | 196.9 | 200.6 | 204.8 | 209.9 |  |
| Average summer temperature_t_ | RSS | 21.7 | **15.72** | 14.73 | 13.96 | 13.74 | 13.37 | 13.07 | 2001 |
|  | BIC | 70.21 | **69.07** | 73.83 | 78.88 | 84.78 | 90.41 | 96.17 |  |
| PC1 | RSS | 64.88 | **44.76** | 37.33 | 33.27 | 30.21 | 30.17 | 35.66 | 2014 |
|  | BIC | 95.4 | **93.13** | 95.22 | 98.85 | 102.9 | 109.14 | 119.25 |  |
| PC2 | RSS | 40.74 | **28.63** | 26.12 | 20.49 | 19.2 | 16.25 | 15.96 | 2000 |
|  | BIC | 84.69 | **82.85** | 87.01 | 87.69 | 92.48 | 94.9 | 100.76 |  |
| PC3 | RSS | 37.47 | 24.96 | **17.85** | 16.09 | 14.36 | 14.25 | 14.18 | 2003, 2006 |
|  | BIC | 82.77 | 79.69 | **78.25** | 82.14 | 85.79 | 91.89 | 98.05 |  |

Table S4

Results of change point analysis on standardized annual abundances of 15 common taxa of arthropods collected in three habitats (arid heath, mesic heath, and fen) at Zackenberg, Greenland between 1996 and 2018. RSS= Residual sum of squares, BIC= Bayesian Information Criterion.

| **Habitat** | **Taxon** |  | **Number of break dates in the model** | | | | | | | **Corresponding break dates** |
| --- | --- | --- | --- | --- | --- | --- | --- | --- | --- | --- |
|  |  |  | **0** | **1** | **2** | **3** | **4** | **5** | **6** |  |
| Arid | Acari | RSS | 2.03 | 1.73 | 1.13 | **0.66** | 0.65 | 0.64 | 0.64 | 2001, 2004, 2014 |
|  |  | BIC | 16.18 | 18.89 | 15.58 | **10.05** | 15.92 | 21.82 | 27.94 |  |
|  | Muscidae | RSS | 1.21 | 0.95 | 0.77 | 0.49 | **0.37** | 0.29 | 0.29 | 1998,2003,2008,2015 |
|  |  | BIC | 3.84 | 4.53 | 5.84 | 1.65 | **1.45** | 2.51 | 8.78 |  |
|  | Chironomidae | RSS | 1.03 | **0.40** | 0.35 | 0.34 | 0.32 | 0.32 | 0.32 | 2015 |
|  |  | BIC | 0.04 | **-15.20** | -12.15 | -6.78 | -1.42 | 4.76 | 11.06 |  |
|  | Coccoidea | RSS | **0.98** | 0.86 | 0.57 | 0.53 | 0.53 | 0.53 | 0.54 |  |
|  |  | BIC | **-1.01** | 2.14 | -0.82 | 3.66 | 9.85 | 15.99 | 22.75 |  |
|  | Collembola | RSS | 2.22 | 0.98 | 0.56 | **0.18** | 0.15 | 0.14 | 0.14 | 1998,2004,2014 |
|  |  | BIC | 17.74 | 5.26 | -1.19 | **-21.52** | -18.59 | -13.78 | -7.52 |  |
|  | Ichneumonidae | RSS | 0.87 | **0.65** | 0.62 | 0.56 | 0.53 | 0.48 | 0.47 | 2015 |
|  |  | BIC | -3.91 | **-4.15** | 0.98 | 4.81 | 10.11 | 13.96 | 19.46 |  |
|  | Linyphiidae | RSS | 0.91 | **0.64** | 0.55 | 0.45 | 0.43 | 0.43 | 0.43 | 2015 |
|  |  | BIC | -2.72 | **-4.68** | -1.68 | 0.04 | 5.34 | 11.37 | 17.64 |  |
|  | Lycosidae | RSS | 1.42 | **0.99** | 0.88 | 0.79 | 0.68 | 0.64 | 0.65 | 1999 |
|  |  | BIC | 7.48 | **5.45** | 9.01 | 12.88 | 15.48 | 20.59 | 27.19 |  |
|  | Sciaridae | RSS | 0.92 | **0.52** | 0.52 | 0.51 | 0.51 | 0.50 | 0.50 | 2015 |
|  |  | BIC | -2.58 | **-9.27** | -3.18 | 2.69 | 8.82 | 15.01 | 21.30 |  |
|  | Nymphalidae | RSS | **1.27** | 1.11 | 0.93 | 0.92 | 0.90 | 0.89 | 0.92 |  |
|  |  | BIC | **4.90** | 8.17 | 10.38 | 16.24 | 22.19 | 28.09 | 35.03 |  |
|  | Phoridae | RSS | 1.29 | 1.02 | **0.31** | 0.27 | 0.25 | 0.23 | 0.23 | 2008, 2011 |
|  |  | BIC | 5.28 | 6.08 | **-15.11** | -12.18 | -7.59 | -2.80 | 3.17 |  |
|  | Thomisidae | RSS | 1.17 | 0.90 | **0.39** | 0.35 | 0.31 | 0.30 | 0.42 | 2005, 2013 |
|  |  | BIC | 2.95 | 3.25 | **-9.61** | -6.14 | -2.08 | 2.89 | 16.97 |  |
| Mesic | Acari | RSS | **1.77** | 1.61 | 1.37 | 1.31 | 1.26 | 1.23 | 1.25 |  |
|  |  | BIC | **13.15** | 17.26 | 19.88 | 25.14 | 30.48 | 36.10 | 42.66 |  |
|  | Muscidae | RSS | **0.72** | 0.60 | 0.54 | 0.50 | 0.45 | 0.44 | 0.43 |  |
|  |  | BIC | **-8.06** | -6.18 | -2.38 | 2.52 | 6.16 | 12.11 | 17.74 |  |
|  | Aphidoidea | RSS | **0.96** | 0.83 | 0.61 | 0.61 | 0.61 | 0.61 | 0.61 |  |
|  |  | BIC | **-1.59** | 1.46 | 0.66 | 6.93 | 13.20 | 19.47 | 25.74 |  |
|  | Chalcidoidea | RSS | 1.79 | 1.27 | 0.49 | 0.34 | **0.23** | 0.22 | 0.27 | 2003, 2006, 2009, 2013 |
|  |  | BIC | 12.82 | 11.20 | -4.50 | -6.27 | **-9.61** | -3.74 | 7.21 |  |
|  | Chironomidae | RSS | **1.89** | 1.50 | 1.28 | 1.16 | 1.13 | 1.13 | 1.13 |  |
|  |  | BIC | **14.02** | 14.96 | 17.65 | 21.69 | 27.31 | 33.57 | 39.84 |  |
|  | Coccoidea | RSS | 1.26 | 0.90 | **0.68** | 0.62 | 0.61 | 0.60 | 0.60 | 2003, 2008 |
|  |  | BIC | 4.67 | 3.35 | **2.97** | 7.21 | 13.11 | 19.06 | 25.33 |  |
|  | Collembola | RSS | 1.02 | **0.47** | 0.36 | 0.33 | 0.31 | 0.29 | 0.28 | 2014 |
|  |  | BIC | -0.18 | **-11.81** | -11.40 | -7.58 | -2.68 | 1.95 | 8.06 |  |
|  | Ichneumonidae | RSS | 0.92 | **0.70** | 0.62 | 0.52 | 0.46 | 0.44 | 0.46 | 1998 |
|  |  | BIC | -2.44 | **-2.47** | 1.06 | 3.22 | 6.83 | 11.77 | 19.09 |  |
|  | Linyphiidae | RSS | **0.84** | 0.72 | 0.64 | 0.58 | 0.52 | 0.51 | 0.50 |  |
|  |  | BIC | **-4.48** | -1.82 | 1.57 | 5.68 | 9.57 | 15.08 | 21.17 |  |
|  | Lycosidae | RSS | 1.67 | 1.16 | **0.56** | 0.52 | 0.45 | 0.42 | 0.48 | 2002, 2013 |
|  |  | BIC | 11.23 | 9.14 | **-1.54** | 2.97 | 6.34 | 10.63 | 20.06 |  |
|  | Sciaridae | RSS | 1.19 | **0.79** | 0.67 | 0.65 | 0.62 | 0.62 | 0.63 | 2014 |
|  |  | BIC | 3.41 | **0.40** | 2.73 | 8.27 | 13.43 | 19.63 | 26.53 |  |
|  | Nymphalidae | RSS | 1.67 | **1.18** | 1.11 | 1.00 | 0.94 | 0.92 | 0.91 | 2008 |
|  |  | BIC | 11.16 | **9.56** | 14.41 | 18.33 | 22.99 | 28.75 | 34.94 |  |
| Fen | Muscidae | RSS | 0.93 | **0.56** | 0.51 | 0.45 | 0.43 | 0.43 | 0.43 | 2004 |
|  |  | BIC | -2.23 | **-7.70** | -3.52 | 0.00 | 5.23 | 11.36 | 17.62 |  |
|  | Chironomidae | RSS | 0.82 | **0.55** | 0.48 | 0.42 | 0.41 | 0.40 | 0.40 | 2014 |
|  |  | BIC | -5.18 | **-7.92** | -4.98 | -1.52 | 3.81 | 9.82 | 15.94 |  |
|  | Collembola | RSS | 1.43 | 1.10 | 0.76 | **0.58** | 0.51 | 0.50 | 0.51 | 1999, 2002, 2014 |
|  |  | BIC | 7.65 | 7.86 | 5.62 | **5.52** | 9.05 | 14.70 | 21.70 |  |
|  | Ichneumonidae | RSS | 1.75 | 1.31 | **0.99** | 0.78 | 0.62 | 0.57 | 0.64 | 2008, 2013 |
|  |  | BIC | 12.26 | 11.98 | **11.71** | 12.46 | 13.46 | 18.03 | 26.87 |  |
|  | Sciaridae | RSS | **2.02** | 1.74 | 1.34 | 1.10 | 1.03 | 1.03 | 1.07 |  |
|  |  | BIC | **15.54** | 18.47 | 18.66 | 20.51 | 25.18 | 31.43 | 38.65 |  |
|  | Scathophagidae | RSS | **1.49** | 1.37 | 1.18 | 1.12 | 1.11 | 1.11 | 1.11 |  |
|  |  | BIC | **8.56** | 12.93 | 15.78 | 20.93 | 26.94 | 33.20 | 39.47 |  |
|  |  | BIC | 11.16 | **9.56** | 14.41 | 18.33 | 22.99 | 28.75 | 34.94 |  |

Table S5

Results of Mann Kendall trend tests of annual standardized abundances of those taxa for which the best model in the change point analysis was identified as being one with no break dates. Significant trends at α = 0.05 are indicated by boldface text.

| Habitat | Taxon | tau | *p* |
| --- | --- | --- | --- |
| Fen | Sciaridae | 0.07 | 0.65 |
|  | Scathophagidae | 0.37 | **0.02** |
| Arid | Coccoidae | 0.14 | 0.40 |
|  | Nymphalidae | -0.01 | 0.96 |
| Mesic | Acari | -0.10 | 0.53 |
|  | Muscidae | 0.06 | 0.74 |
|  | Aphidoidae | -0.44 | **0.01** |
|  | Chironomidae | -0.24 | 0.13 |
|  | Linyphiidae | -0.30 | 0.06 |
